# Wearable facemask-attached disposable printed sensor arrays for point-of-need monitoring of ammonia in breath

**DOI:** 10.1101/2024.07.16.603629

**Authors:** Giandrin Barandun, Abdulkadir Sanli, Chun Lin Yap, Alexander Silva Pinto Collins, Max Grell, Michael Kasimatis, Jeremy B. Levy, Firat Güder

## Abstract

Blood sampling, despite its historical significance in clinical diagnostics, poses challenges such as invasiveness, infection risks, and limited temporal fidelity for continuous monitoring. In contrast, exhaled breath offers a non-invasive, pain-free, and continuous sampling method, carrying biochemical information through volatile compounds like ammonia (NH3). NH3 in exhaled breath, influenced by kidney function, emerges as a promising biomarker for renal health assessment, particularly in resource-limited settings lacking extensive healthcare infrastructure. Current analytical methods for breath ammonia, though effective, often face practical limitations. In this work, we introduce a low-cost, internet-connected, paper-based wearable device for measuring exhaled ammonia, designed for early detection of kidney dysfunction at the point-of-need. The device, which attaches to disposable facemasks, utilizes a disposable paper-based sensor array housed in a biodegradable plastic enclosure to mitigate high relative humidity (RH) issues in breath analysis. We validated our technology using a laboratory setup and human subjects who consumed ammonium chloride-containing candy to simulate elevated breath ammonia. Our wearable sensor offers a promising solution for rapid, point-of-need kidney dysfunction screening, particularly valuable in resource-limited settings. This approach has potential applications beyond kidney health monitoring, including chemical industry safety and environmental sensing, paving the way for accessible, continuous health monitoring.

## Introduction

Blood has historically been the sample matrix of choice when searching for or measuring biochemical markers of health and disease in the body.^[1]^ Accessing blood for biochemical analysis, however, is challenging for at least three reasons: i) Blood samples require painful procedures for extraction (this is especially problematic in children); ii) the risk of infection is increased when the skin barrier is damaged and; iii) for an extensive range of measurements, the amount of blood needed limits the frequency of analysis, therefore reducing temporal fidelity that requires continuous-time measurements, reducing diagnostic performance.

The respiratory rate of a healthy human is between 12-18 breaths times per minute.^[2]^ Unlike blood, breathing provides easy access to a gaseous sample matrix (that is, exhaled breath) carrying information concerning the internal biochemistry of the body through the exchange of gases in the respiratory tract, including the mouth.^[3]^ Breath-based, non-invasive, pain-free assessments can be performed continuously over time with a high frequency of measurement, which is not easily achievable through blood-based measurements with implants or skin-attached microneedles.^[4]^

Human breath contains a range of volatile organic and inorganic compounds, including biomarkers such as isoprene, ammonia, and acetone, which can be used to predict various disease states.^[5–7]^ Ammonia (NH_3_) is a toxic byproduct of protein metabolism, which is produced microbially in the gut and cellularly throughout the body.^[8]^ NH_3_ and its ionized form NH_4_^+^ are removed from the body either directly or by conversion into urea (CH₄N₂O) in the liver through the urea cycle.^[9]^ Urea is a highly water-soluble, practically non-toxic, small molecule that is stable in the body.^[10]^ Urea can be rapidly hydrolyzed into CO_2_ and NH_3_ catalytically in the presence of urease, an enzyme that is only produced microbially; hence, urease is not an endogenous enzyme.^[10]^ Because of its small size, urea diffuses readily into tissues, including the oral cavity and saliva.^[10]^ When kidneys function abnormally, urea concentrations in the blood and tissues increase; therefore, blood urea concentration (BUN) is used as a diagnostic biomarker to assess kidney health.^[11]^ In the oral cavity, urea is hydrolyzed by the urease-positive microbes into NH_3,_ which is expelled from the body with exhaled breath. ^[12]^

Patients suffering from end-stage kidney failure from any cause exhale higher ammonia levels in their breaths (820 ppb – 14700 ppb, with a mean of 4880 ppb) compared to healthy individuals (425 – 1800 ppb, mean of 960 ppb).^[13]^ Kidney dysfunction, caused by chronic kidney disease (CKD) or acute kidney injury (AKI), is currently diagnosed by measuring serum creatinine (sCreat) and BUN, both blood-based tests.^[14]^ Assessing kidney health non-invasively, rapidly, and at a low cost is especially important in resource-limited settings (RLS) where blood testing is not widely available, such as in middle- and low-income countries.^[15]^ In developing nations where healthcare is severely lacking outside major centers, AKI that goes undiagnosed due to lack of diagnostic testing leads to preventable deaths, and CKD can only be diagnosed currently by blood testing, causes no symptoms, is an increasing world-wide public health issue, and potentially treatable.^[16]^ Exhaled NH_3_ is, therefore, particularly suited for use as a diagnostic marker to measure kidney (dys)function in RLS to prevent premature deaths due to CKD and AKI; an estimated 13.3 million cases of AKI occur in developing nations annually.^[17]^

To avoid blood-based testing, to date, analytical techniques, such as gas chromatography/mass spectrometry (GC-MS), selected-ion flow-tube mass spectrometry (SIFT-MS), laser spectroscopy, and laser photoacoustic spectroscopy, have been developed for measuring breath ammonia.^[18–22]^ These methods are, however, bulky, costly, and impractical for use in RLS as a diagnostic technique due to pre-analytical errors and challenges associated with sample handling. Other approaches, such as quartz crystal microbalance^[23]^, chemical^[24]^, and optical sensors^[25]^ can perform with high precision at trace levels, but they face limitations in terms of analytical performance. For most sensing technologies, the high content of moisture in breath (human breath is > 90% in relative humidity (RH)) creates a myriad of analytical issues.^[26]^ Moisture can poison catalytic surfaces and block sensing surfaces by adsorption or condensing into liquid droplets, which in turn damage sensors or electronics.^[27]^ The presence of high content of water in breath, therefore, limits the analytical performance of most low-cost sensing approaches. High RH of exhaled breath has been the primary factor preventing the development of breath-based diagnostics.^[28,29]^

In this work, we report a low-cost, internet-connected, paper-based wearable device for measuring exhaled ammonia in human breath, with the goal of producing a non-invasive method for early detection of kidney dysfunction at the point-of-need (**Figure 1**). The technology reported consists of a disposable paper-based sensor array to overcome RH-related artifacts, housed in a biodegradable plastic enclosure. The device attaches to disposable facemasks commonly used in healthcare. The test results are transmitted to a nearby smartphone for post-processing and can be shared with a remote professional over the internet; hence, they are compatible with telemedicine.^[30]^ We validated our approach through ammonia using a characterization environment in our laboratory and human subjects, who consumed a candy (salty licorice) that contains NH_4_Cl to generate NH_3_ in exhaled breath.

**Figure 1:**
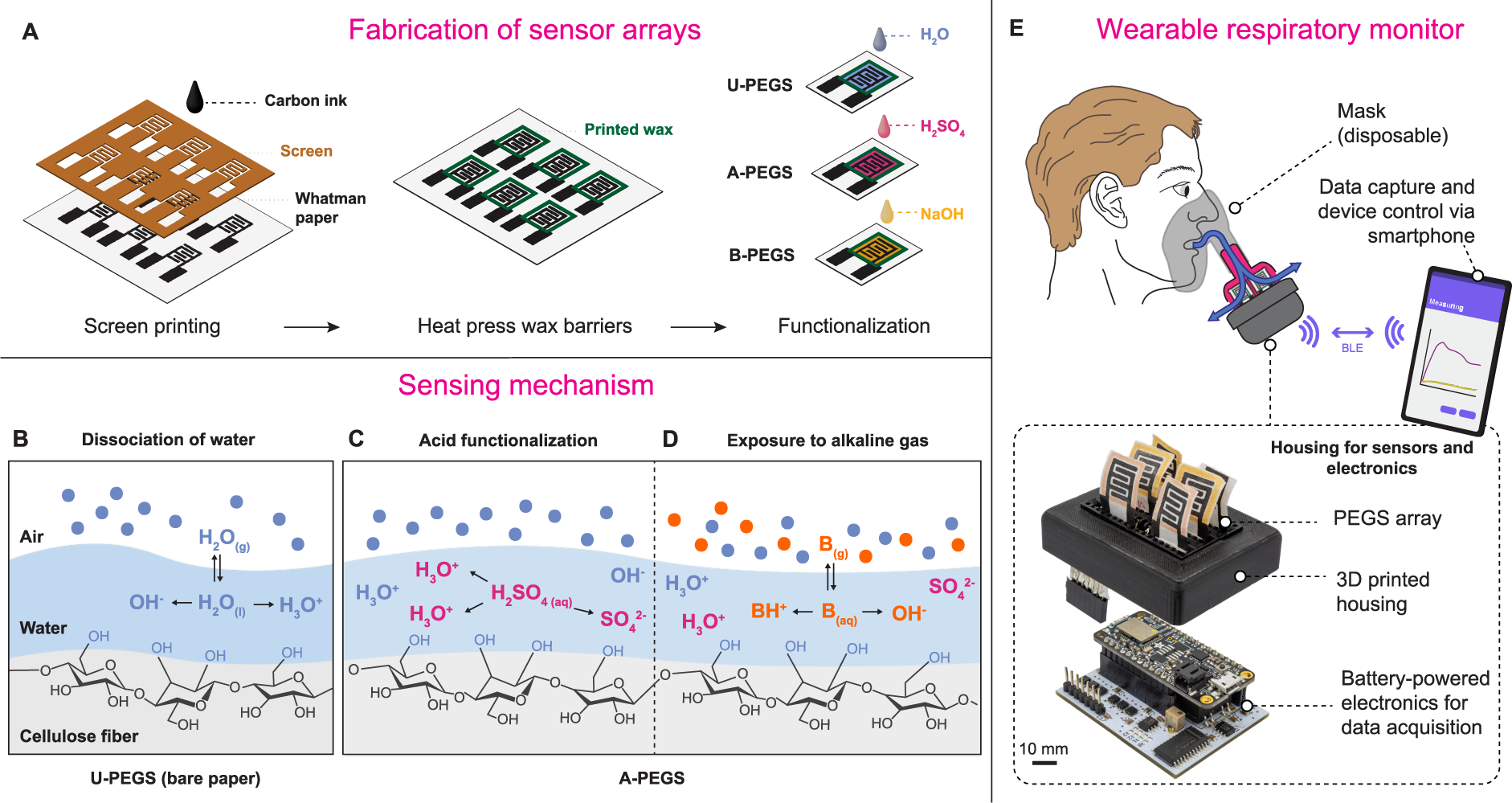
Fabrication of sensor arrays. **(A)** Illustration of carbon ink electrodes on chromatography paper with wax squares for liquid confinement. **(B)** Mechanism of conductivity in the paper due to water accumulation around cellulose fibers with increasing relative humidity (RH). **(C)** Functionalization process involving sulfuric acid (H_2_SO_4_) on paper electrodes to modify electrical impedance. **(D)** Response of acid-treated sensors to alkaline gases such as ammonia, affecting electrical impedance. **(E)** Application of sensor arrays in a wearable respiratory monitor attached to a medical mask, with Bluetooth low energy (BLE) connectivity for data transmission. Illustrations not to scale.

## Results and Discussion

### Fabrication of sensors

Paper-based gas sensors (PEGS) are produced by screen printing carbon electrodes onto Whatman^TM^ chromatography paper, allowing rapid prototyping and low-cost ($0.02/sensor) production of PEGS (**Figure 1A**). For selectively detecting NH_3_ in exhaled breath, we used an array of sensors that consisted of three types of functionalization: (i) Untreated-PEGS (U-PEGS), produced by adding 10 µL of deionized (DI) water; (ii) base-treated PEGS (B-PEGS), produced by adding 10 µL of sodium hydroxide (0.001 M – 0.1 M NaOH) and; (iii) acid-treated PEGS (A-PEGS), produced by adding 10 µL of sulfuric acid (0.001 M – 0.1M H_2_SO_4_) to the paper. Wax barriers created around the carbon electrodes prevented the spread of the solutions of acid or base, spatially confining the functionalized regions of paper. Because the chemical compounds added slowly react with environmental gases such as CO_2_, the functionalization was performed right before the start of the experiments, no longer than 15 minutes before. Notably, NH₃ has the highest concentration in exhaled breath among alkaline gases, and PEGS produces the highest response to NH₃ among all gases tested; ^[31]^ Therefore, in this work, NH₃ is the primary alkaline gas we are sensing.

### Mechanism of Sensing

Paper is a highly hygroscopic material that consists of natural, microfibers of cellulose, which absorb moisture from the immediate environment. At higher levels of RH (> 40%), the water present around the fibers of cellulose within paper behaves like bulk water.^[32]^ Ions in the layer of water adsorbed in the paper can move freely through the network of cellulose fibers, rendering paper electrically more conductive (**Figure 1B**).^[31]^ A water-soluble gas that dissolves and dissociates into ions in water can, therefore increase the number of ions in paper, leading to higher electrical conductivity.^[31]^ The electrical conductivity (σ) of water depends on the concentration (n_ion_), charge (Z_e_), and mobility (µ_ion_) of ions present such that σ = n_ion_ × Z_e_ × µ_ion_. When paper is pre-treated with H_2_SO_4_, two additional hydronium ions (2H_3_O^+^) and one sulfate ion (SO_4_^2-^) are produced for each H_2_SO_4_ molecule dissolved in water (**Figure 1C**), leading to higher electrical conductivity. When, however, an alkaline gas, such as ammonia (NH₃) reacts with the sulfuric acid functionalized paper (A-PEGS), ammonia (NH₃) dissolves in the water layer on the cellulose fibers and dissociates to form water (H₂O), hydroxide ions (OH⁻), and ammonium ions (NH₄⁺). The hydroxide ions (OH⁻) neutralize the acidity from the sulfuric acid, resulting in the formation of water (H₂O) and ammonium sulfate ([NH₄]₂SO₄). [NH₄]₂SO₄ is a highly water-soluble salt and will be present in its dissociated form (two ammonium ions (2NH₄⁺) and one sulfate ion (SO₄²⁻)) when dissolved. The addition of NH₃ to aqueous sulfuric acid substitutes hydronium ions (H₃O⁺) with ammonium ions (NH₄⁺). (**Figure 1D**). Because NH_4_^+^ ions have a lower mobility than H_3_O^+^ (7.63×10^-4^ vs 36.23×10^-4^ cm^2^ V^-1^s^-1^), the reaction of NH_3_ with A-PEGS functionalized with H_2_SO_4_ causes a drop in the electrical conductivity of paper.^[33]^ The mechanism of sensing acidic gases with B-PEGS is similar to A-PEGS, which increases selectivity toward the detection of acidic gases.

The electrical conductivity of paper changes both when it reacts with a water-soluble gas or when the RH increases. A single PEGS (with or without chemical modifications) would, therefore, have low selectivity when operating in a multi-component mixture of gas. To increase selectivity toward a target acidic or alkaline gas (i.e. NH_3_) in the presence of fluctuating levels or RH, we used an array of sensors consisting of U-PEGS, A-PEGS, and B-PEGS. The responses produced by each sensor within the array could then be used for differential analysis to calculate the concentration of the target gas as different sensors would react with the target gas to a varying extent. Integrated into a disposable facemask, the array of sensors would allow non-invasive monitoring of levels of exhaled breath ammonia where the RH would be changing in each cycle of inhalation and exhalation (**Figure 1E**).

### Characterization

To study the behavior arrays of sensors consisting of U-PEGS and A-PEGS and U-PEGS and B-PEGS, we exposed the arrays to different concentrations of NH_3_ and CO_2_ (relevant gasses for the analysis of levels of NH_3_ in exhaled breath) while keeping the RH at 65% (**Figure 2**). Fixing the RH constant enabled us to characterize the behavior of each sensor (array) toward the target analytes in a more precise fashion, important for understanding the underlying phenomena. To account for the intrinsic microstructural variability of paper and chemical modifications, the electrical responses originating from each sensor was normalized to the baseline conductance – i.e. ΔG / G_0_ (G_0_ is the electrical conductance of PEGS in the absence of the target gas; ΔG is the change in electrical conductance when exposed to the target gas).

**Figure 2:**
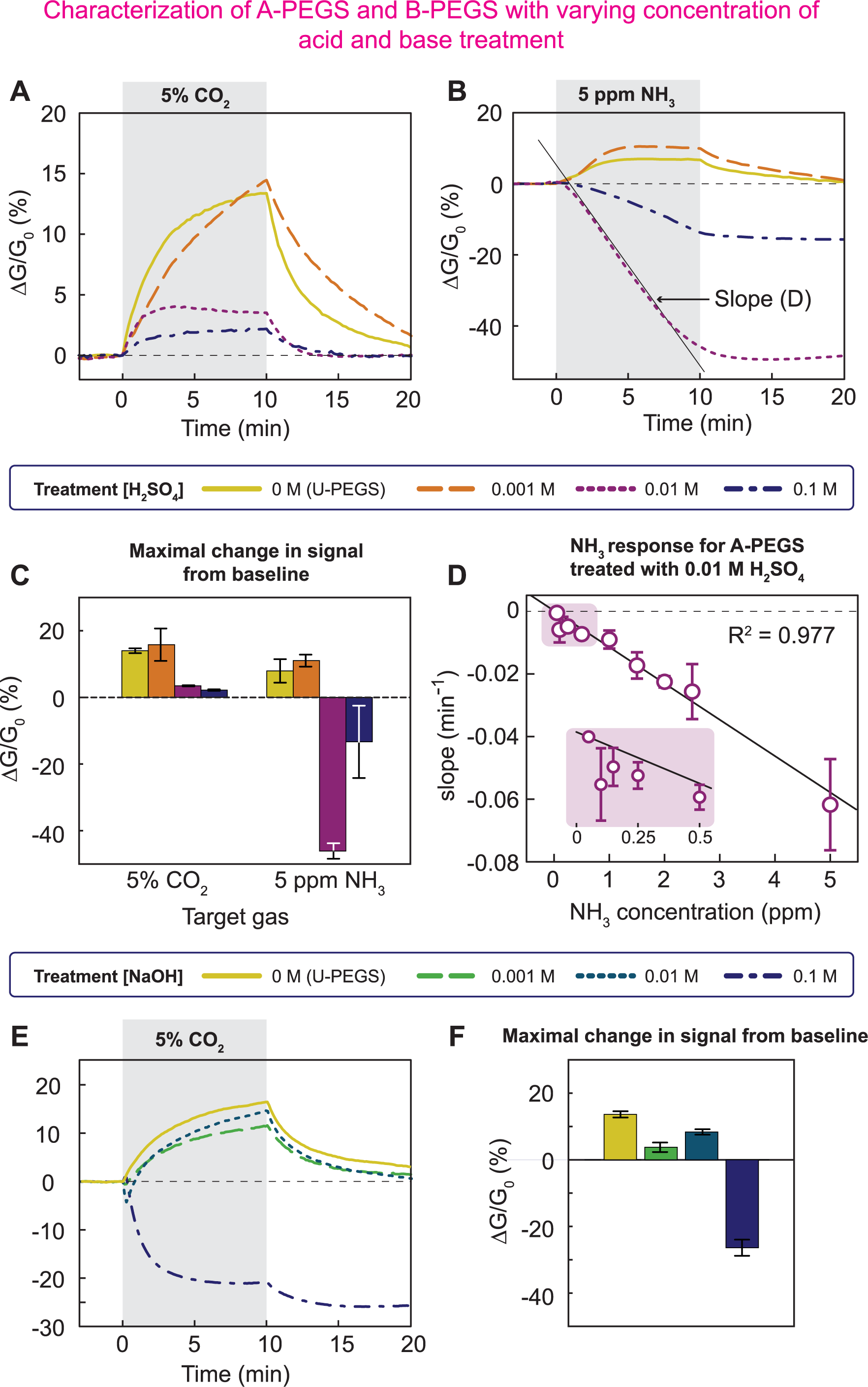
Characterization of A-PEGS and B-PEGS with varying concentration of acid and base treatment in the test chamber. The conductance of PEGS is measured when a sinusoidal signal of 4 V and 10 Hz is applied to the sensors (see **SI-P3** for details). (**A**) Four PEGS were treated differently: one U-PEGS and three A-PEGS with different concentrations of H_2_SO_4_. All PEGS were exposed to 5% CO_2_ for 10 min. The bar plot shows maximal signal changes in a 10 min interval for the differently treated sensors, and the error bars indicate the standard deviation for n=3. (**B**) The same array used in (A) is exposed to 5 ppm of NH_3_ for 10 min. For high acid concentrations (0.01 M and 0.1 M), the alkaline gas neutralizes the acidic pre-treatment, and the signal drops constantly over time (slope is the black solid line). For the lowest concentration (0.001 M), the acid is depleted quickly, and the sensor starts behaving like a U-PEGS. (**C**) Maximal change in signal from baseline for the A-PEGS exposed to 5% CO_2_ and 5 ppm of NH_3_. (**D**) A-PEGS treated with 0.01 M H_2_SO_4_ shows the best signal in terms of error and especially sensitivity to NH_3_ (see (C)). We exposed these A-PEGS to a wide range of NH_3_ concentrations (0.05 ppm to 5 ppm) and calculated the slope of the drop (black solid line in (B)). This gives a linear correlation between slope and NH_3_ concentration with a coefficient of determination of R^2^ = 0.977. At lower concentrations (pink inlet), the errors get bigger. B-PEGS characterization with varying concentrations of alkaline treatment (**E**) Four PEGS were treated differently: one U-PEGS and three B-PEGS with different concentrations of NaOH. One bare-PEGS is compared to three B-PEGS when exposed to 5% CO_2_ for 10 min. (**F**) The acidic gas neutralizes the alkaline pre-treatment and the signal for B-PEGS initially drops. For low concentrations of base (0.001 M and 0.01 M) the PEGS are depleted quickly, and the signal starts increasing again. The error bars are the standard deviation for n=3-12.

When exposed to 5% CO_2_, A-PEGS (treated with 10 µL of 0.001 M to 0.1 M H_2_SO_4_) shows a reversible response (**Figure 2A**). We observed that increased concentration of H_2_SO_4_ decreases the sensitivity of CO_2_ in two ways: (i) The base conductance (G_0_) is higher with sensors containing higher amounts of H_2_SO_4_, therefore and the relative change (ΔG) is smaller; (ii) Decreasing pH reduces the solubility of CO_2_ in water leading to lower increases in net ionic strength hence lower sensitivity.^[34]^

When the array was exposed to 5 ppm NH_3,_ we observed two different behaviors (**Figure 2B**). For the lowest concentration of H_2_SO_4_ (0.001M, orange, long dashed), A-PEGS behave like U-PEGS and show an increase in response. For higher concentrations of added sulfuric acid (0.01 M, 0.1 M), we observed a continuous drop in signal. This behavior can be explained by ongoing neutralization of the H_2_SO_4_ by the dissolved NH_3_ in paper. When the concentration of H_2_SO_4_ was small, initial H_2_SO_4_ was immediately neutralized without much effect on the overall electrical conductance. When the concentration of the acid increased, exposure to NH_3_ produced a noticeable drop in the response of the A-PEGS. The rate of the change of conductance over time (slope) could, therefore, be used as an indicator of the NH_3_ concentration (e.g., straight grey line in Figure 2B for 0.01M H_2_SO_4_). Comparing the treatment with 0.01M H_2_SO_4_ to 0.1M (**Figure 2C)**, we found that the sensitivity to 5 ppm NH_3_ is approximately three times higher, and the standard deviation is five times smaller (n=3) for A-PEGS treated with 0.01M H_2_SO_4_.

**Figure 2D** shows the relative rates of change of conductance (min^-1^, straight grey line Figure 2B) for A-PEGS treated with 0.01M H_2_SO_4_ when exposed to concentrations of NH_3_ ranging from 0.05 to 5 ppm (see **Figure S1**). We observed a linear correlation between the slope of the response of A-PEGS and NH_3_ concentration (R^2^ = 0.977) as exposure to higher levels of NH_3_ neutralizes the acid more quickly. For lower concentrations (0.05-0.25 ppm), the data exhibited higher standard deviation and lower linear correlation (R^2^_0.05-0.25ppm_= −0.95). Longer exposure times improve the limit of quantification (LOQ) because the increased total amount of NH_3_ passing the sensors enhances detection sensitivity.

For B-PEGS, the response to CO_2_ is more complicated (**Figure 2E**). For 0.1M NaOH (purple, dot-dashed) we see a drop in conductance as the base gets neutralized by the acidic gas. After gas exposure is stopped, we see another drop in conductance because additional bicarbonate ions (HCO_3_^-^) form CO_2_, which is released from the sensor into the environment again. For a lower base concentration (0.01M NaOH, green, long dashed), the base is neutralized within the first minute (sharp drop in conductance). After that, bicarbonate ions form in the sensing element, which increases conductance. For the lowest base concentration (0.001M NaOH, cyan, dashed), we do not see a neutralization effect. The base gets neutralized by the acidic gas within a few seconds and the sensor shows a conductance increase due to bicarbonate ions forming in the sensing element.

### Detecting exhaled NH_3_ in a respiratory simulator

In each cycle of breathing, the RH immediately outside the oronasal opening fluctuates between 100% RH and room RH (inhalation). To characterize the performance of the PEGS array in the presence of fluctuating levels of RH and NH_3_, we built a respiratory simulator (**Figure S2**). The simulator cycles the RH in the gas sensor characterization chamber between 100% and 45% RH (the RH in our laboratory) at a (adjustable) frequency of six breaths per minute while introducing a controlled concentration of NH_3_ in each cycle of exhalation.

We first subjected a PEGS array consisting of three A-PEGS and three U-PEGS to simulated cycles of respiratory activity without any NH_3_ (**Figure 3A**). The desorption of moisture from paper is thermodynamically less favorable than adsorption.^[35]^ Simulated cycling of respiratory activity, therefore, slowly increases the moisture content within paper, eventually reaching a steady state after 7-8 min regardless of the amount of acid added to paper in the context of A-PEGS. Next, we introduced 5 ppm of NH_3_ along with respiratory cycling (**Figure 3B**). Although NH_3_ started neutralizing H_2_SO_4_ present in A-PEGS immediately, we first observed a steady increase in the response of the sensor. After about 4 minutes, the A-PEGS with 0.01M H_2_SO_4_ exhibited a steady drop in response as expected. The initial increase and subsequent decrease in the response of the sensor can be attributed to two competing processes moisture build-up *vs* neutralization of H_2_SO_4_ in which the latter eventually dominates the overall electrical conductance. All the other sensors showed primarily an increase in the response of the sensor since adsorption of moisture dominated the electrical signal.

**Figure 3:**
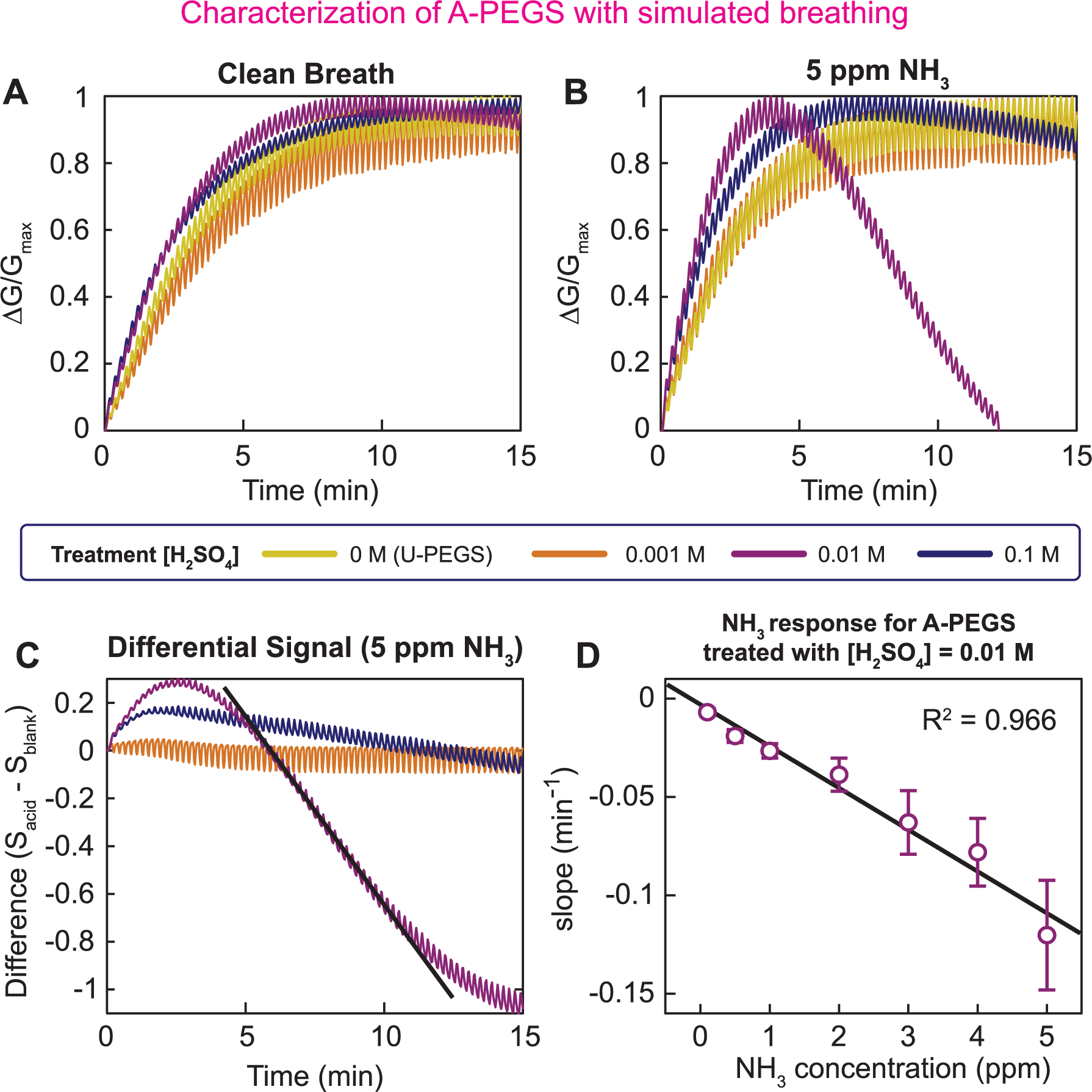
Characterization of A-PEGS with simulated breathing. We tested PEGS arrays consisting of U-PEGS and A-PEGS in our respiratory simulation chamber. Breathing was simulated by exposing the PEGS in turns to dry air (ca. 50% RH) and humidified air (ca. 90% RH). We mixed different concentrations of NH_3_ (0.1 ppm – 5 ppm) into the compressed air to simulate ammonia in breath. (**A**) The response of the array to clean breath (no ammonia). The signal rises until the sensors reach an equilibrium with the breathing. We normalized the signal for better comparison and all sensors behave similarly. (**B**) We exposed the array to breath containing 5 ppm NH_3_. The U-PEGS and lowest acid concentration A-PEGS (0.001M) show no change to clean breath. We see a decline after the initial peak for the A-PEGS treated with higher concentrations of H_2_SO_4_ (0.01M and 0.1M). Similar to the previous tests, the A-PEGS with 0.01M H_2_SO_4_ shows the highest sensitivity and the decline can be seen clearly. **(C)** To decouple the signal from any environmental influence (especially RH), we take the difference in the signal of the U-PEGS and the A-PEGS (S_A-PEGS_ – S_U-PEGS_). The slope of the differential signal after the initial peak (dashed line) can now be used to determine the NH_3_ concentration. (**D**) We plotted the slope (straight black line in (C)) for A-PEGS treated with 0.01M H_2_SO_4_ against different concentrations of NH_3_ in breath ranging from 0.1 ppm to 5 ppm. The data shows a linear correlation (R^2^ = 0.966). The error bars indicate the standard deviation for n=3.

We exploited the dominance of the RH response as opposed to neutralization in the non-or slightly modified sensors in the array for differential analysis (**Figure 3C**). Differential analysis yields a curve in which the effect of moisture is subtracted from NH_3_ thereby allowing calculation of the drop hence NH_3_ concentration in exhaled (simulated) breath. Mathematically, the isolation of the alkaline gases, such as NH_3_ signal is achieved by subtracting the signal of the U-PEGS, which primarily responds to RH changes from the signal of the A-PEGS, which responds to both NH_3_ and RH. Here, the equation used is S_NH3_=S_A-PEGS_-S_U-PEGS_, where S_U-PEGS_ is the total response of the A-PEGS, and S_U-PEGS_ is the response of U-PEGS to RH. The slope of the differential signal measured at varying levels of NH_3_ produced a linear relationship (R^2^ = 0.966) with a limit of detection (LOD) is 0.1 ppm of NH_3_ after 15 minutes (**Figure 3D**) simulated respiratory cycling. To achieve lower LOD than 0.1 ppm a longer duration of respiratory cycling may be necessary.

### Human testing

In a series of experiments, we tested our approach for measuring levels of NH_3_ in exhaled breath, we produced a wireless sensor module (see **SI-P3**) that can be attached to a disposable facemask for wearable, point-of-care analysis of breath (**Figure 1E**). The PEGS array used in these experiments comprised three A-PEGS treated with 0.01M H_2_SO_4_ and three U-PEGS. We tested our respiratory device with eight healthy male volunteers from our research group (body mass index (BMI) 18-26, age 22-34). Because all subjects participating in the study were healthy, to simulate kidney disease, we asked the volunteers to consume salty licorice candy (Malaco Salmiak Balk Sweet from *Scandinavian Candy & Sweets*) which contains copious amounts of NH_4_Cl. The human experiments (**Figure 4)** consisted of two parts. (i) We first asked the volunteers to wear the respiratory monitor for 15 minutes and breathe normally through their mouths; (ii) we then gave each volunteer a piece of candy and once again asked them to wear and breathe through the mask while consuming the candy without chewing.

**Figure 4:**
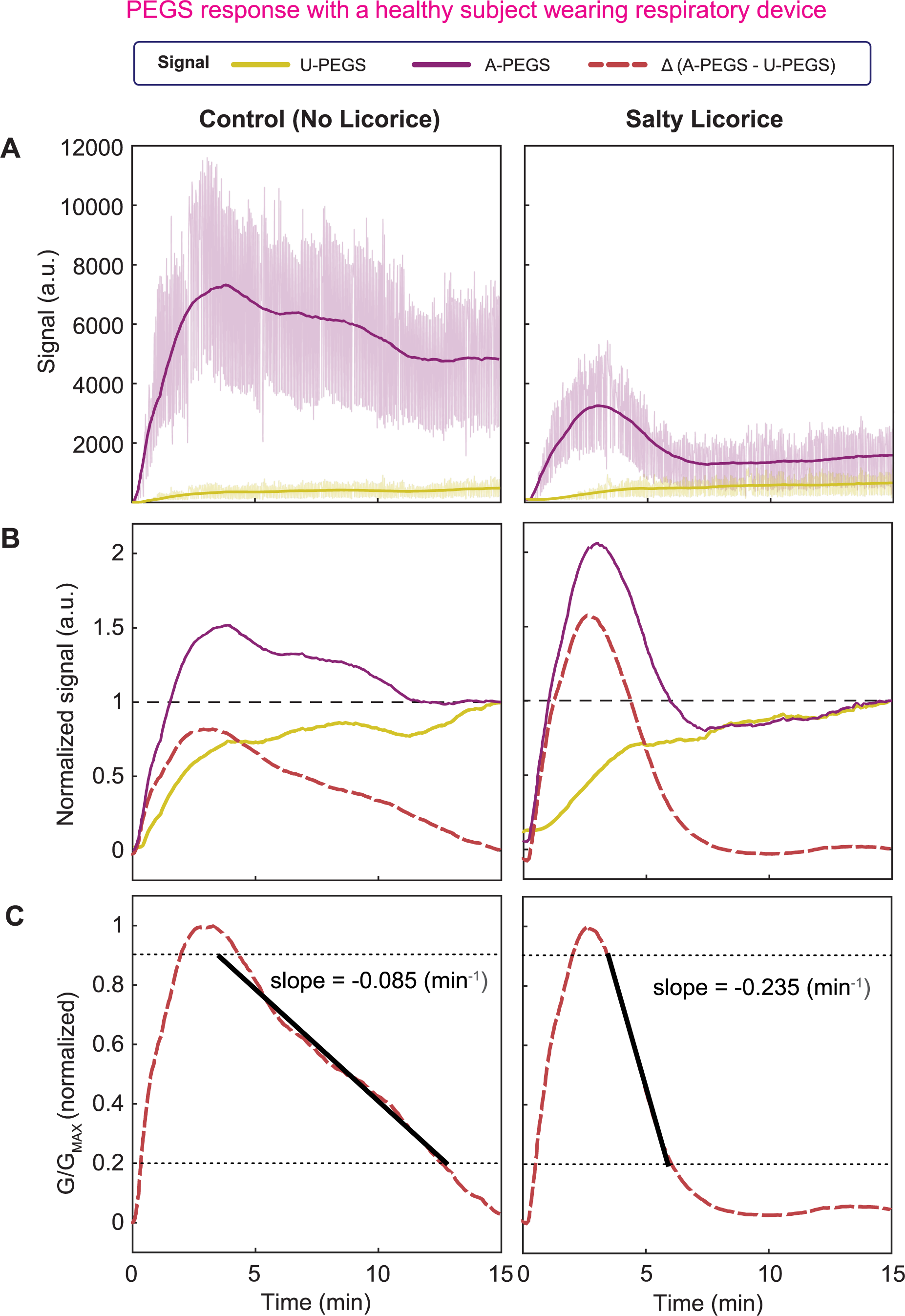
PEGS response with a healthy subject wearing respiratory device. The raw signal obtained from a human subject breathing into the facemask, therefore on the sensors, for 15 min is shown in green for A-PEGS and blue for U-PEGS. To eliminate the signal fluctuations due to inhalation and exhalation, we use a 1000-point moving average (pink line). This can track the conductivity changes in the PEGS from humidity and ion changes in the breath without the fluctuations of breathing. The human subjects were asked to do a control experiment (normal breathing, left) and an experiment where they were eating a salty licorice while doing the respiratory experiment (right). (**B**) We divide the moving average from (A) by the end point after 15 min to relate the signal from U-PEGS and A-PEGS. Then, we subtract the two signals to filter the changes from an increased RH. This difference tracks the drop of the A-PEGS due to the neutralization of the sulfuric acid by alkaline gases (i.e. NH_3_). (**C**) We normalized the difference from Figure 4B to compare different subjects and experiments. The slope is calculated on an interval from 90% peak height to 20%. The straight grey dashed lines show the interval we use to calculate the slope of the drop. Control experiments with no salty licorice (left) show a flatter decrease than experiments with salty licorice (right).

For differential analysis, we first applied a 1000-point averaging low-pass filter to smoothen the signals acquired from the sensors (**Figure 4A**) as the raw measurements were noisy. Next, we normalized each signal to the final data point acquired at the 15-minute mark (**Figure 4B**) to allow subtraction of the response generated by U-PEGS from A-PEGS to produce a differential signal. We finally calculated the slope of the differential signal in the linear region before the signal flatlined which would indicate the completion of the neutralization reaction between exhaled NH_3_ and H_2_SO_4_ present in paper.

Our tests (**Figure 5A**) with the healthy volunteers (n=8) showed a statistically significant difference (paired t-test; *p* <0.05) between the healthy and control groups (i.e. simulated diseased state) when the subjects consumed either no candy or a single full candy. The slope in the control experiment might come from small amounts of ammonia that can be present in healthy humans.^[36]^ Even though the amount of ammonia might vary from subject to subject, the concentration in healthy human breath is negligible compared to the concentration in breath following the consumption of salty licorice. We conducted another experiment with a single volunteer who was asked to consume smaller quantities of the candy (n=3): ½ and ¼ of a single candy (**Figure 5B**). Although there was a clear difference between the average slopes calculated for the control (no candy), ½, and ¼ of a single candy experiment, increasing the candy amount from ¼ to ½ does not make a significant difference according to the statistical tests (one-tailed t-test).

**Figure 5:**
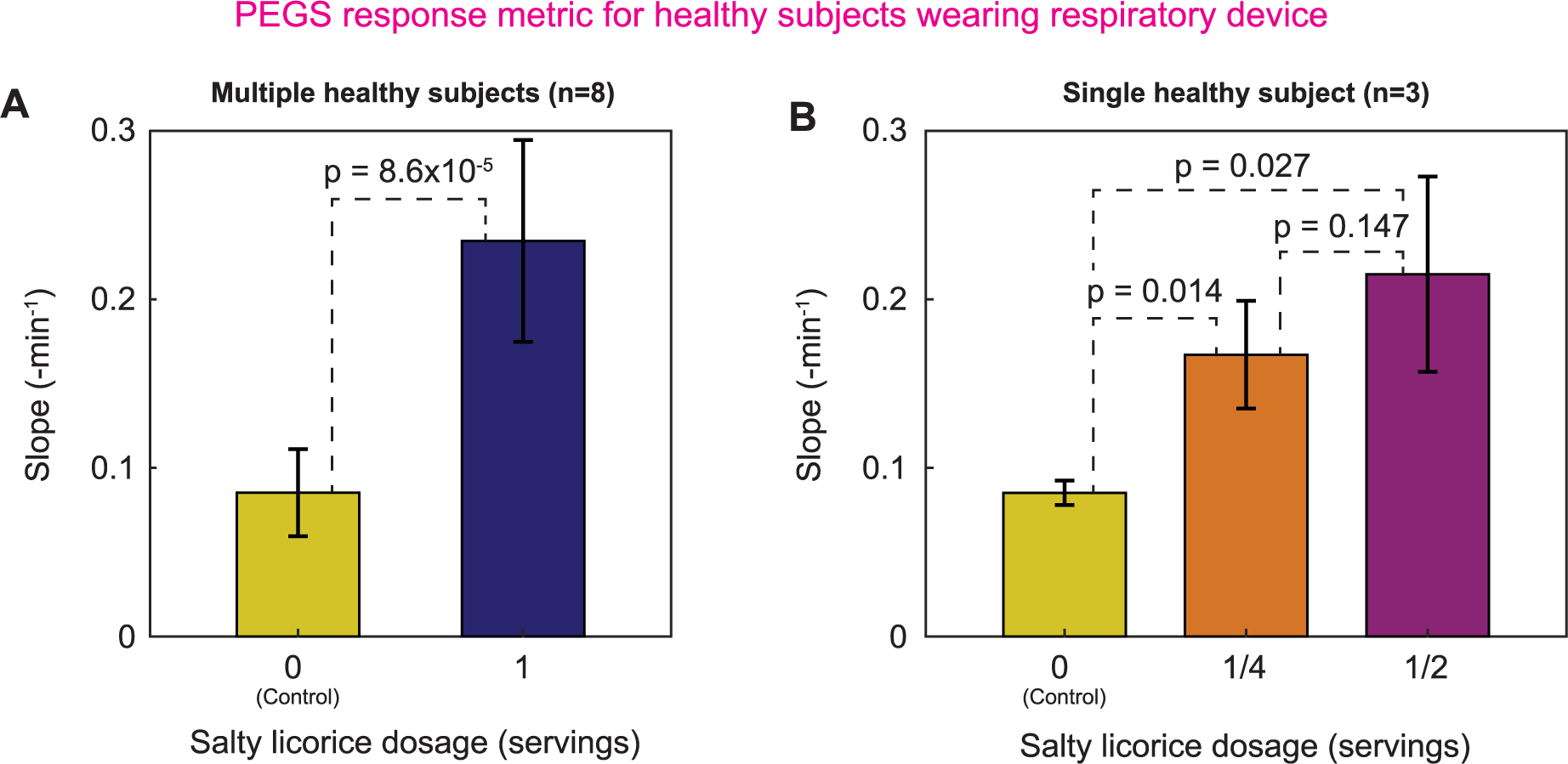
(**A**) The results of the mean slope (see Figure 4C) for eight healthy subjects in an experiment with normal breathing (control) and breathing while eating a salty licorice. The slope increases three-fold when the ammonia from the candy neutralizes the sulfuric acid in the A-PEGS. (**B**) A similar experiment to (A) with one subject only and three repeats on each bar. In addition to the control experiment, the subject was asked to eat ¼ and ½ of a salty licorice during the 15 min experiment. The error bars indicate the standard deviation for (A) *n=8* and (B) *n=3*. The *p*-values come from one-tailed (right) paired samples *t*-test, indicating a significant difference if *p* < 0.05.

## Conclusion

The sensing technology reported is a low-cost approach to analyze the chemical composition of exhaled breath across a large range of concentrations (i.e. high dynamic range) without depending on collecting a breath condensate which is the most common method of analyzing exhaled breath^[37,38]^ (see **Table S2**). By not depending on collecting condensation, it is possible to rapidly detect chemical compounds with simpler instrumentation. For our laboratory prototypes, each sensor costed approximately US $0.02 to produce which would need to be replaced before each measurement. The electronics and plastic housing costed US $75 (see **Table S1**) however these components are reusable after disinfection. All the components of the plastic housing were made of biodegradable polymers and can also be disposed of which would of course slightly raise the unit price of each test. Although in this work, the plastic components were 3D printed, they can also be injection molded hence every single element within our design is compatible with the existing high-volume manufacturing methods except for wax printing which would need to be replaced with an appropriate alternative. There is, therefore, a large scope for reducing the cost of the device proposed potentially one to two orders of magnitude at large scales of production.

The sensing platform reported, however, has at least four disadvantages: (i) The screen-printed graphite electrodes are susceptible to cracking if the paper is creased, but this is unlikely during use as the sensors were securely placed inside the respiratory device without folds. (ii) Performance of PEGS is hindered by the lower levels of relative humidity (< 20%) and temperatures below 0 °C due to the freezing point of water. Additionally, because paper is an organic substrate, PEGS are unsuitable for operation at high temperatures (> ∼120 °C) though such conditions are rare for biological applications. (iii) The impedance of PEGS ranges from kΩ to GΩ, limiting the scope of simplification for the electronics. (iv) The arrays of PEGS also face a limitation in miniaturization due to the inherently larger size of paper-based structures compared to silicon more sophisticated, microfabricated devices. Nevertheless, given the wearable, standardized formfactor (facemask) of our approach, further miniaturization would not substantially improve use or costs.

Using the PEGS array, we were clearly able to detect the consumption of licorice candy that contains ammonium chloride by eight healthy human subjects. We were, however, not able to detect the amount of candy (dose, from ¼ to ½) in a statistically significant fashion although clearly the averages were visibly different. We believe that the way the candy is consumed (chewed vs slowly sucked) impacts the measurement results of the exhaled analyte (NH_3_). The human experiments can be improved by more standardization of the testing protocols and consumption of licorice can be a simple model to simulate diseased states similar to CKD or AKI. The performance of the sensor array can also be further improved in two different ways: i) by performing measurements for a longer time for detecting low levels of analyte or; ii) by increasing the pH of the oral cavity which would in turn improve the release of gaseous NH_3_.

Although, in this work, we primarily focused on the use of PEGS array for detecting NH_3_ in human breath, its potential applications extend far beyond the topic described in this study. PEGS array can be utilized in various fields such as the chemical industry for monitoring hazardous gases, medical diagnostics, agriculture, and environmental monitoring.^[27,39–41]^ Because our approach uses a mobile device for data collection, the data produced by the PEGS array can be easily processed and stored on the cloud, enabling remote access by healthcare professionals which is especially important in Low-or Middle-Income Country (LMIC) with limited access to care. If mobile operation is not required, a simple LCD display can also be implemented into our design to provide information at the point of care without connectivity. In the future, we will validate the application of PEGS arrays in clinical experiments to measure the levels of exhaled ammonia in patients suffering from AKI and CKD to monitor BUN levels non-invasively.

## Materials and Methods

### Wearable respiratory monitor

For field testing, we developed a wearable respiratory monitor easily attached to a commercially available disposable medical facemask (**Figure 1E**). The respiratory monitor for human testing consists of a medical facemask, plastic housing for PEGS array (sensor chamber), and electronics (**Figure S3**). The sensor chamber contains six PEGS: three U-PEGS and three A-PEGS. The electronics chamber provides the electronic circuit to read, process, and transmit the sensor data. We designed the read-out component in-house and used Bluetooth low energy (BLE) to read and control the device with a smartphone (see **SI-P3**). Human test subjects are asked to wear the whole setup and breathe normally through the mouth for 15 min. The exhaled breath passes the PEGS array and leaves through holes at the bottom of the sensor chamber.

Based on the guidelines at Imperial College London, the project team performed a detailed risk assessment to identify the hazards and risks associated with non-invasive human experiments. All human subjects gave consent for the sensor testing experiments. All experiments were conducted on healthy subjects.

### Experimental setup for breathing test

We conducted four experiments in the following two different environments: in the first, a laboratory environment, to characterize the PEGS arrays, we designed a test chamber with known RH, flow rate, and gas concentration (see **SI-P2**). We can control the test environment and its parameters with three different gas lines: humid compressed air, which we humidified by passing it over the headspace of a DI water container; dry compressed air; and the target gas (i.e. NH_3_ or CO_2_) in air. In the second, we monitored healthy individuals with our wearable respiratory monitor (**Figure 1 FiguC**).

Our experiments consisted of two characterization experiments in the sensor chamber and two tests on human subjects: in the first experiment, we exposed the PEGS array in controlled RH and flow rate to the target gas (NH_3_ and CO_2_) in our sensor chamber. We applied simulated breath (see below) with a tidal volume of ca. 270 ml and a breathing rate of 6 breaths/min. Both tidal volume and breathing rate are smaller than in average humans (tidal volume: 7 ml/kg, breathing rate: 10-20 breaths/min) ^[42]^ and are optimized for our system. We checked the PEGS array response to different concentrations of NH_3_ from 0.05 ppm to 5 ppm in simulated exhalation. With our wearable respiratory monitor, we measured the NH_3_ content in the breath of healthy individuals before and after eating salty licorice (*Malaco Salmiak Balk Sweet* from Scandinavian Candy & Sweets). This Scandinavian candy contains a quantity (ca. 4-8%) of ammonium chloride (NH_4_Cl). While sucking on the candy, NH_3_ forms in the saliva and mixes into the exhaled breath through the mouth.

### Simulated human breathing

When breathing, humans do not use the full capacity of the lungs. If no extra effort is applied, the air exhaled while breathing is typically called tidal volume. An average human has a tidal volume of ca. 7 ml/kg and breathes between 10-20 times/min.^[42]^

To simulate human breathing, we programmed mass flow controllers (MFCs, type GM50A from MKS) in our test set-up. The process involves a humidified air flow (>90% RH, 2000 ml/min) passing over the sensors for 8 seconds, representing exhalation. This is followed by a 4-second airflow with relative humidity similar to the room (approximately 50% RH, 2000 ml/min), representing inhalation. We found that these timings closely matched the patterns of human breathing. The simulation, however, is not perfect and has the following shortcomings: i) The exhaled volume is twice the inhaled volume. ii) The tidal volume in this setup is approximately 270 ml, half of the average human tidal volume (ca. 500 ml). iii) The simulation completes 6 cycles per minute, slower than the average 8-12 breaths/min in humans. iv) The temperature is constant, unlike the temperature difference between body and room temperature in human respiration. *v*) We do not add CO_2_ to the exhalation, using the same cylinder of compressed air for both processes. Despite the mentioned shortcomings, our test set-up can simulate the general shape of a breathing curve over time. To further mimic human respiration, we added different ammonia concentrations (0.1 ppm – 5 ppm) to the exhalation stream, simulating oral ammonia production in humans.

## Acknowledgments

The authors would like to thank the Department of Bioengineering at Imperial College London. F.G. would like to thank the Bill and Melinda Gates Foundation (Grand Challenges Explorations scheme under grant numbers OPP1212574 and INV-038695). A.S. would like to thank the Bill and Melinda Gates Foundation (INV-038695). F.G and A.S acknowledge Innovate UK BMC (Reference: PA2131). F.G. would like to thank the US Army (U.S. Army Foreign Technology (and Science) Assessment Support (FTAS) program under grant number: W911QY-20-R-0022) for their generous support. F.G. and G.B. would like to thank EPSRC (EP/R010242/1), General Electric Healthcare and Imperial College, Department of Bioengineering for their generous support. F.G, and A.S.P.C. thank EPSRC (EP/L016702/1) and BBSRC DTP (Reference: 2177734). M.K. acknowledges EPSRC DTP (Reference: 1846144). F.G and M.G. thank the Imperial College Centre for Plastic Electronics and acknowledge EPSRC for Plastic Electronics Doctoral Training Centre (EP/G037515/1 and EP/L016702/1). J.L. is supported by the National Institute for Health Research (NIHR) Biomedical Research Centre based at Imperial College Healthcare NHS Trust and Imperial College London.

