## Supplementary Information for "Wearable facemask-attached disposable printed sensor arrays for point-of-need monitoring of ammonia in breath"

### Table of Contents

SI-P1. Characterization of A-PEGS for different $NH_3$ concentrations
SI-P2. Test chamber for PEGS characterization
SI-P3. Electronics and software
SI-P4. Comparable technologies in literature

### SI-P1. Characterization of A-PEGS for different $NH_3$ concentrations

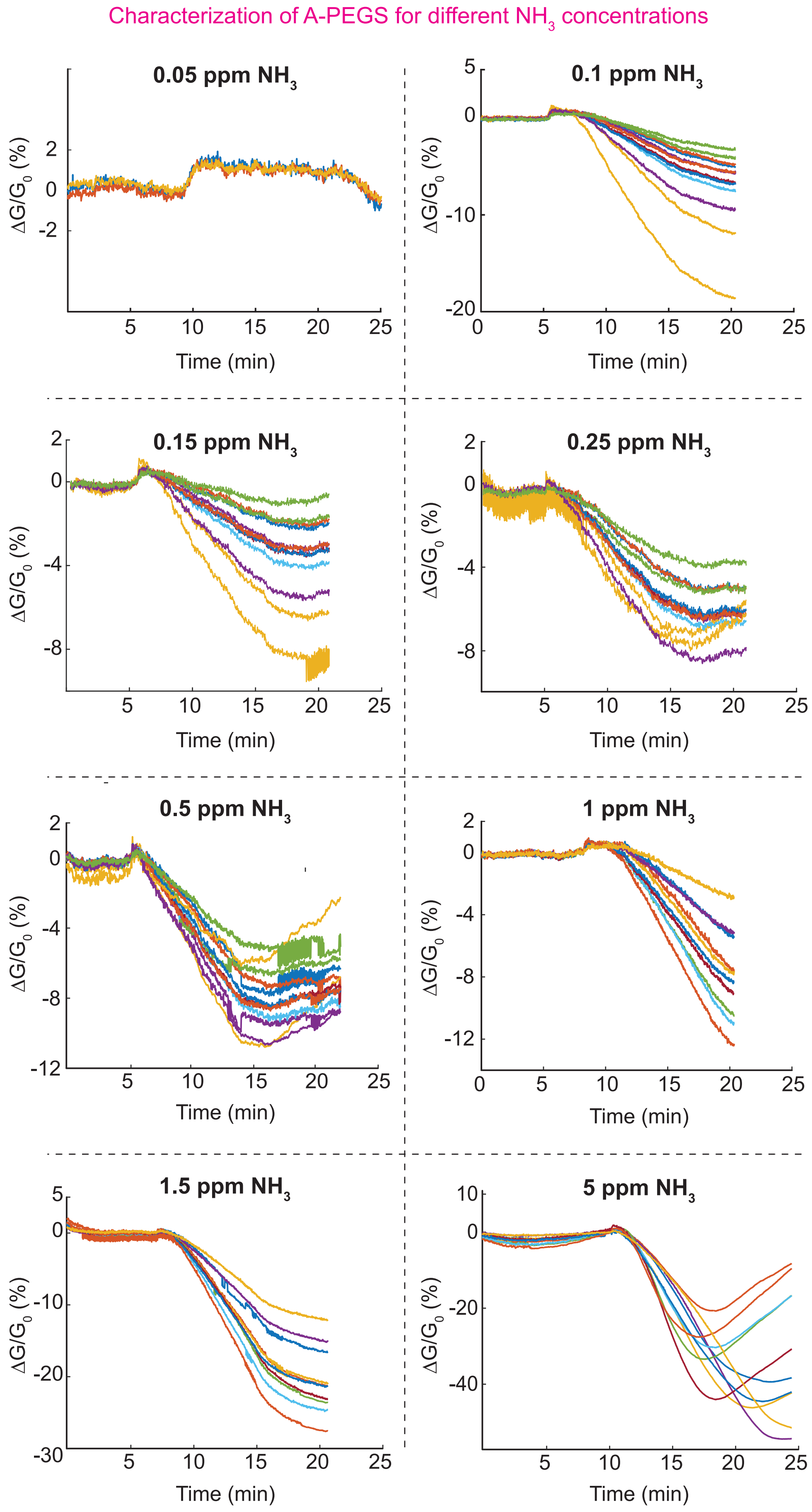

**Figure S1**: Characterization of A-PEGS. Change of conductance of A-PEGS for different amounts of $NH_3$ from 0.05 ppm to 5 ppm. With the increase of ammonia concentration, the slope also increases, indicating a linear correlation between $NH_3$ concentration and the conductance slope. n=8-12.

**Figure S1** shows the raw signal when exposed to ammonia in time ($min^{-1}$) is shown for different concentrations of ammonia from 0.05 ppm to 5 ppm. For low concentrations (0.05 ppm to 0.25 ppm) the data shows higher standard deviation and less reliability on the linear correlation. We reach our lower limit of detection (LOD) at ca. 0.1 ppm for our test set-up. The current set-up, however, is limited in terms of mixing, flow rate and gas concentration. For low concentrations of ammonia, we mix high flow rates of compressed air (2000 ml/min) with very low flow rates of ammonia (10 ml/min). If the gas mixture is not perfectly homogeneous the target gas might show spatial differences in concentration (i.e. some areas of the chamber have higher concentrations whereas other areas are not reached at all. This can explain the higher errors and difficulties in detecting.

### SI-P1. Test chamber for PEGS characterization

Three mass flow controllers (MFCs) (type GM50A from Bronkhorst UK Ltd) are programmed to adjust the flow rates of the three streams to keep the parameters in the test chamber at certain levels and can be controlled from a computer (**Figure S2**). The two lines of carrier gas are used to reach a precise RH level in the supply stream. This stream is mixed with the target gas and supplied into the test chamber. The test chamber is a polytetrafluoroethylene (PTFE) box (120×40×60 $mm^3$) with two inlets at the top and one inlet on each small side to provide an evenly distributed supply of gas.

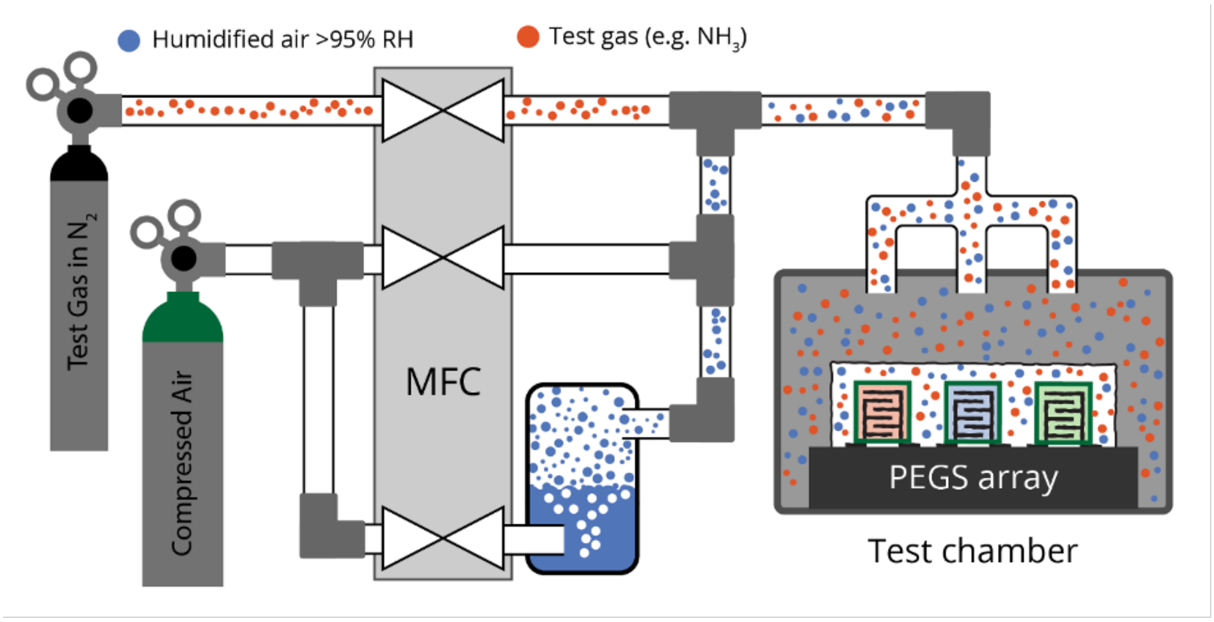


**Figure S2**: Test chamber set-up for the sensor characterization experiment. The carrier gas (compressed air) can either be used dry or humidified. To humidify the carrier gas, it bubbles through DI water (blue circles). The test gas (red circles) is mixed with the dry and humidified carrier gas. The mixing ratio is controllable using MFCs to achieve the desired RH and test gas concentration inside the test chamber containing the sensors.

With this set-up, we can test for a gas over a wide range of concentrations at different RH levels. For example, for ammonia we have a range of 1% to 100 ppb (part per billion). The flow rate of each line was adjusted with MFCs with a total flow rate of 2000 mL/min reaching the test chamber. The current set up, however, is limited in terms of mixing, flow rate and gas concentration. For low concentrations of ammonia, we mix high flow rates of compressed air (2000 ml/min) with very low flow rates of ammonia (10 ml/min). If the mixing is not perfect the target gas will reach areas of the chamber quicker or not reach other areas at all. This can explain the higher errors and difficulties in detecting.

We preferred intersurgical face masks available at EcoLite™ due to their widespread availability and affordability. We believe, however, that other brands could also serve the purpose effectively. Furthermore, we developed a 3D printed polylactic acid (PLA) housing for electronics to enhance the functionality of the masks.

### SI-P2. Electronics and software

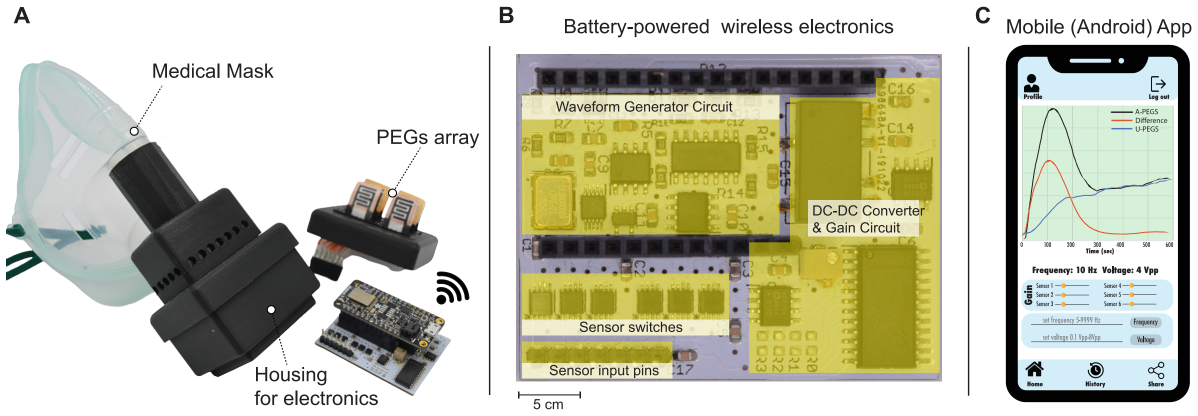


**Figure S3**: **Integrated system overview**. (**A**) Exploded view of the mask housing PEGS array and electronics for measurements. (**B**) Photograph of the battery-powered wireless electronics for sensor data acquisition. (**C**) Image of the mobile (Android) app displaying and analyzing individual PEGS array readings, enabling comprehensive data visualization and analysis. The Android app was designed and implemented in the Android Studio development environment using Java and XML programming languages. We used an android phone (model Huawei P30 Lite) for our testing.

**Figure S3** provides an integrated system overview. In (**A**), an exploded view illustrates the medical mask housing the PEGS array and associated electronics for measurements. We preferred intersurgical face masks available at EcoLite™ due to their widespread availability and affordability. We believe, however, that other brands could also serve the purpose effectively. Furthermore, we developed a 3D printed polylactic acid (PLA) housing for electronics to enhance the functionality of the masks. Figure S3B shows a photograph of the battery-powered wireless electronics used for sensor data acquisition. **Figure S3C** displays an image of the mobile Android app, which visualizes and analyzes individual PEGS array readings for comprehensive data analysis. The app was developed in the Android Studio environment using Java and XML programming languages, and we used a Huawei P30 Lite phone for testing.

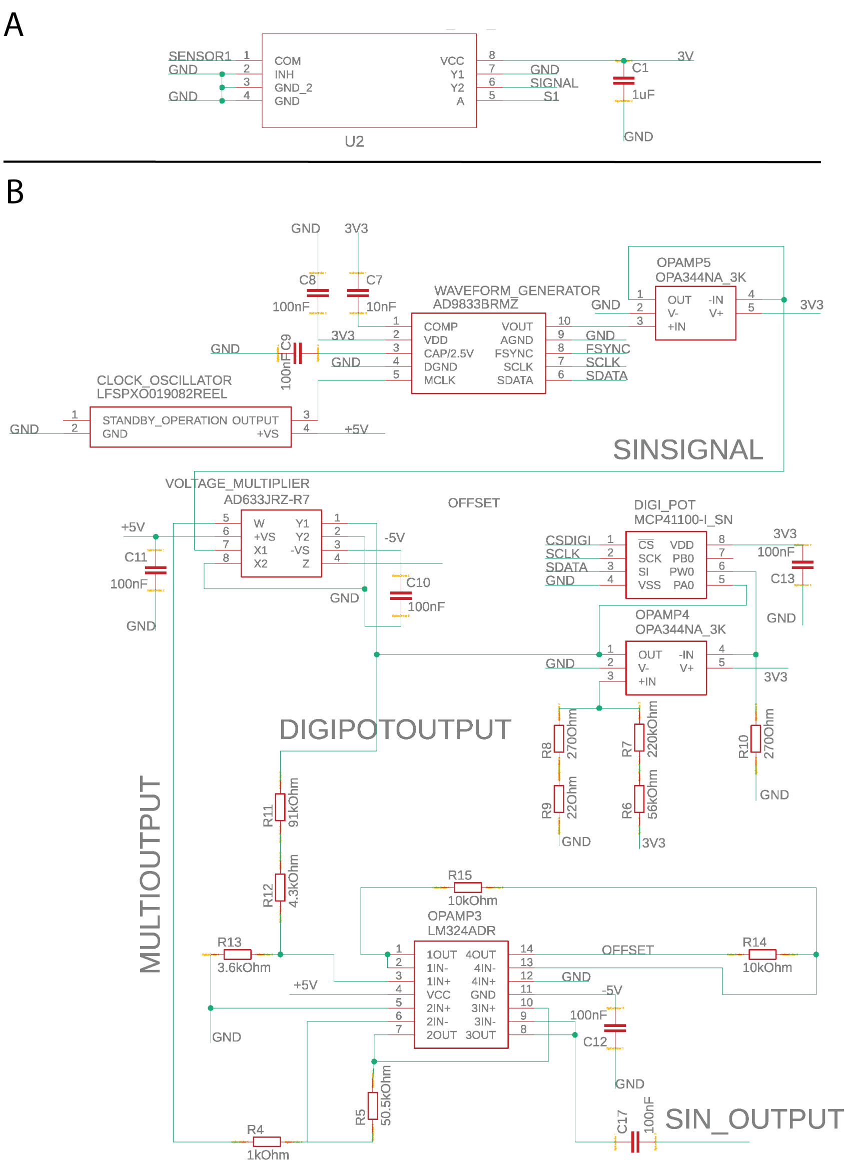


**Figure S4:** **(A)** Analog switch circuit for PEGS, **(B)** This circuit creates the sinusoidal wave to measure impedance in the PEGS array. A 24MHz clock oscillator clocks a waveform generator (AD9833) which is fed into a voltage follower (OPAMP5) to create the base signal 'SINSIGNAL' in a frequency between 10Hz-10kHz. The DC offset of the sinusoidal signal is then removed using a voltage multiplier (AD633) and amplified in a transimpedance amplifier configuration using an adjustable gain (digital potentiometer, MCP41100) to reach the desired voltage range (max. ca. +/- 4.2V). The final signal 'SIN_OUTPUT' is the input to the PEGS.

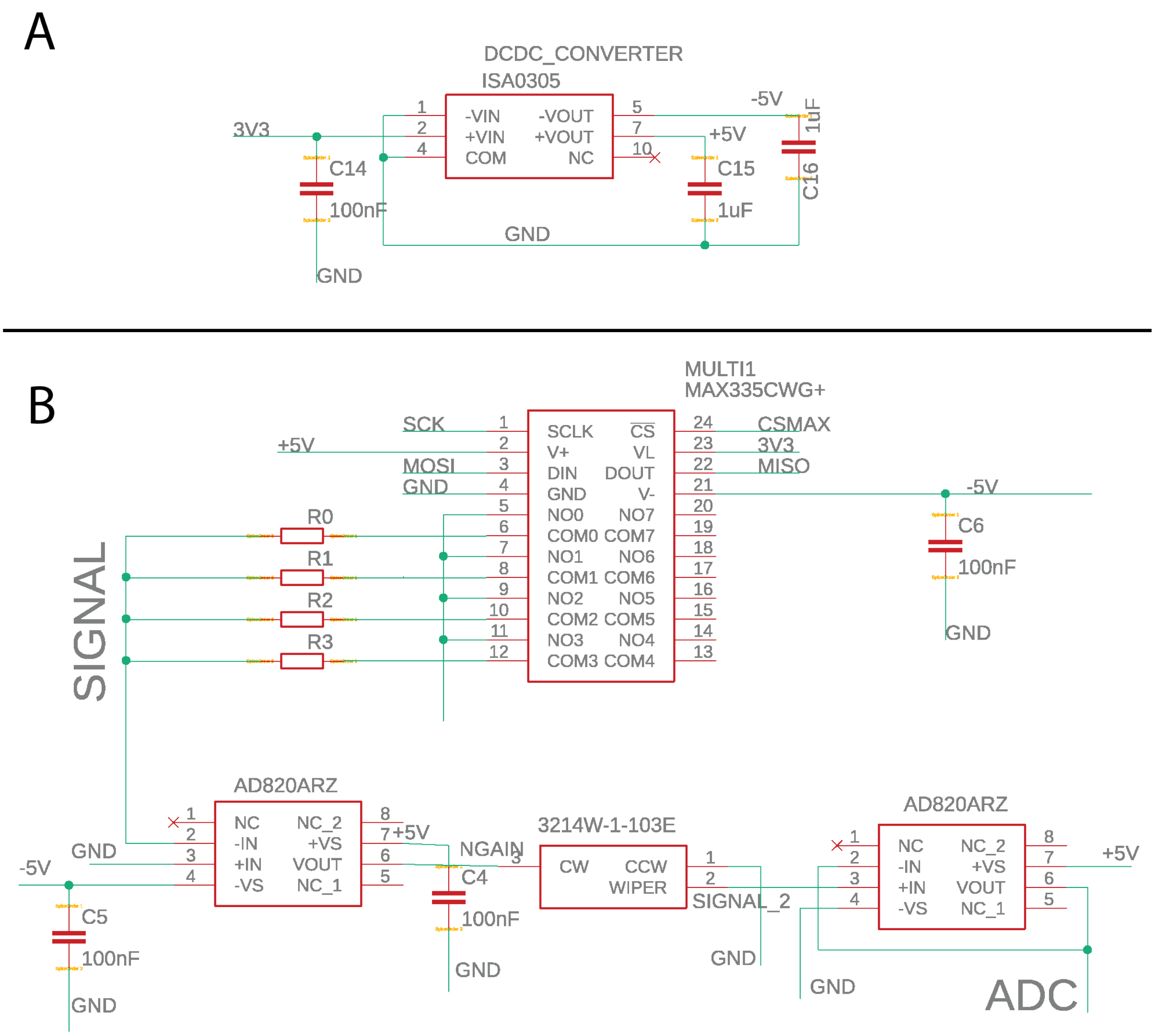


**Figure S5**: (**A**) A DC-DC converter is used to extend the 3.3V from a standard Arduino to -5 / +5 V to achieve proper AC sinusoidal waveforms (see Figure 11). **(B)** This circuit takes the sensor signal as an input, amplifies, and feeds the signal into the Arduino ADC. The sensor input is ‘SIN_OUTPUT from Figure S3 and the output from the sensor (‘SIGNAL’) is amplified using a transimpedance amplifier configuration. A multiplexer (MAX335) chooses the correct gain resistor (50M, 10M, 1M, 100k) and a potentiometer (3214W) is used to downscale the 4Vpp signal range to 3.3V to be read by the Arduino ADC.

To measure the response to target gases, a sinusoidal excitation with an amplitude of 4 Vp-p at 10 Hz was applied, and conductance (G) was determined through Ohm’s law, enabling the calculation of $\Delta G/G_0$ to assess the sensor's reaction to the gas. For the impedance measurements, we followed the methodology outlined in our previous paper [3] (see specifically Figure 3). This choice was based on considerations of the Debye-Falkenhagen effect, and the capacitive charging behavior observed at different frequencies.

**Table S1**: Prices of components (in USD) used for the fabrication of the electronic circuits and the corresponding equivalent platform using Bluetooth for data communication.

| **Component** | **Price* (USD)** |
|---|---|
| Bluefruit LE Development Board (Adafruit Feather M0) | 24.95 |
| Waveform Generator (AD9833) | 11.87 |
| Analog Voltage Multiplier (AD633JRZ-R7) | 12.44 |
| Analog switches (MAX335) | 7.29 |
| DC-DC Converter (ISA0305) | 3.70 |
| Precision Operational Amplifier (AD820ARZ) | 6.38 |
| Resistors/ Capacitors/ Potentiometer | 2.15 |
| PCB manufacturing | 4.69 |
| PLA Housing | 0.67 |
| Li-Po Battery charger circuit | 1.49 |
| **TOTAL (USD)** | **75.63** |

* Prices per unit are based on ordering 10 units and are converted from GBP using the prevailing exchange rate, subject to change. All components were purchased from Mouser Electronics.

### SI-P3. Comparable technologies in literature

**Table S2**: 1) A device tested in real-time on a human subject; 2) Humidity controllable test chamber but not mentioned at what RH experiments are conducted; 3) DFB-QCL: Distributed feedback quantum cascade laser (not handheld); 4) QCM: Quartz crystal microbalance sensor; 5) Additional equipment to control the humidity is needed for real-time breath analysis; 6) ssDNA-FG: single-stranded DNA-functionalized graphene; 7) TFB: (poly[(9,9-dioctyl-fluor-enyl-2,7-diyl)-co-(4,4′-(N-(4-s-butylphenyl)diphenylamine)]); 8) IL-SOWG: Ionic liquid-based slab optical waveguide sensor; 9) D-A: electron donating and electron accepting; 10) PVP: poly(vinyl pyrrolidone), impedimetric; 11) NP: nanoparticles; 12) Exhaled breath was passed over a quicklime bag and additionally, corrections to the signal due to humidity changes were calculated (humidity sensor needed).

| Technology/ Material | Humidity | LOD | Real-time1) | Year [Ref] |
|---|---|---|---|---|
| **PANI nanojunction** | dried | 16 ppb | sample bag | 2008[4] |
| **$MoO_3$** | controlled2) | 50 ppb | simulation | 2010[5] |
| **DFB-QCL3)** | real breath | 6 ppb | yes | 2011[6] |
| **PANI nanoparticles** | real breath | 40 ppb | yes | 2013[7] |
| **QCM4) ($SiO_2$)** | constant | 1000 ppb | samples | 2015[8] |
| **Si-doped α-$MoO_3$** | constant (90%) | 400 ppb | no | 2015[9] |
| **CuBr** | constant (40%) | 10 ppb | yes5) | 2016[10] |
| **ssDNA-FG6)** | constant (80%) | 103 ppb | no | 2017[11] |
| **TFB7)** | dried (10%) | <100 ppb | sample bag | 2017[12] |
| **IL-SOWG8)** | dried | 69 ppb | sample bag | 2018[13] |
| **D-A9) polymer nanopores** | dried (10%) | 100 ppb | sample bag | 2019[14] |
| **PVP10)** | constant (97%) | 500 ppb | sample chamber | 2019[15] |
| **Au NP11)-$V_2O_5$/$CuWO_4$** | constant (n.a.) | 212 ppb | no | 2020[4,16] |
| **CuBr film** | dried12) | 100 ppb | yes | 2020[4] |
| **$BaFe_{12}O_{19}$ NP11)** | dried | 200 ppb | no | 2020[17] |
| **PEGS Array (this work)** | real breath | 100 ppb | yes | 2024 |
